# Probabilistic Genotyping of Single Cell Replicates from Complex DNA Mixtures Recovers Higher Contributor LRs than Standard Analysis

**DOI:** 10.1101/2021.11.05.467485

**Authors:** Kaitlin Huffman, Erin Hanson, Jack Ballantyne

## Abstract

DNA mixtures are a common source of crime scene evidence and are often one of the more difficult sources of biological evidence to interpret. With the implementation of probabilistic genotyping (PG), mixture analysis has been revolutionized allowing previously unresolvable mixed profiles to be analyzed and probative genotype information from contributors to be recovered. However, due to allele overlap, artifacts, or low-level minor contributors, genotype information loss inevitably occurs. In order to reduce the potential loss of significant DNA information from donors in complex mixtures, an alternative approach is to physically separate individual cells from mixtures prior to performing DNA typing thus obtaining single source profiles from contributors. In the present work, a simplified micromanipulation technique combined with enhanced single-cell DNA typing was used to collect one or few cells, referred to as direct single-cell subsampling (DSCS). Using this approach, single and 2-cell subsamples were collected from 2-6 person mixtures. Single-cell subsamples resulted in single source DNA profiles while the 2-cell subsamples returned either single source DNA profiles or new mini-mixtures that are less complex than the original mixture due to the presence of fewer contributors. PG (STRmix™) was implemented, after appropriate validation, to analyze the original bulk mixtures, single source cell subsamples, and the 2-cell mini mixture subsamples from the original 2-6-person mixtures. PG further allowed replicate analysis to be employed which, in many instances, resulted in a significant gain of genotype information such that the returned donor likelihood ratios (LRs) were comparable to that seen in their single source reference profiles (i.e., the reciprocal of their random match probabilities). In every mixture, the DSCS approach gave improved results for each donor compared to standard bulk mixture analysis. With the 5- and 6- person complex mixtures, DSCS recovered highly probative LRs (> 10^20^) from donors that had returned non-probative LRs (<10^3^) by standard methods.

## 1. Introduction

DNA mixtures are a commonly encountered source of evidence found at crime scenes and are one of the more difficult sources of biological evidence to interpret due to their inherent complexity [1] as multiple genotype combinations can often explain the STR mixture data. This is particularly troublesome since much DNA evidence is comprised of a mixture originating from multiple contributors including the victim (s) and/or perpetrator(s) to the crime or to others who have no direct relationship to the incident in question [2–4]. With the implementation of probabilistic genotyping (PG), mixture deconvolution analysis has been revolutionized as PG is less subjective, more consistent, uses more DNA profile information, and provides greater statistical power resulting in the analysis of mixtures that were previously unresolvable [5,6].

Even with the implementation of PG software, probative information from all contributors to a mixture is not always achieved [5] due to a loss of genotype information caused by overlapping alleles, artifacts such as stutter, or very low-level minor contributors [7–10]. Furthermore, if the number of contributors (NOC) to a mixture is underestimated, information about some of the donors cannot be inferred [3]. Other factors such as the relatedness of the donors [11] or > 4 donors contributing to the mixture, can lead to even more complexity in allele overlap resulting in further uncertainty of NOC. Even if the correct NOC is identified, not all donors will result in a probative LR value. Moreover, if a reference profile from a person of interest (POI) is not available, analysis is severely limited as PG analysis alone does not truly deconvolute mixtures into their single source genotypes in most instances, but rather provides the likelihood (in the form of a likelihood ratio) that the DNA results indicate a specific genotype could have contributed to the mixture at hand. Without a reference genotype to condition the LR on, or an appropriate database to search against, actionable probative information cannot always be obtained. This is particularly true of complex mixtures which can be defined as samples containing comingled DNA from ≥ 2 individuals where stochastic effects or allele sharing leads to uncertainty in determining each of the contributors’ DNA genotypes [12]. An exception to this deconvolution issue would be the analysis of a simple 2-person mixture with clear distinction of the major and minor contributor in which there were no significant stochastic and low-template effects [13].

A potential solution to overcoming the issue of genotype information loss is to separate individual cells or a small subset of cells from the original ‘bulk’ mixture prior to analysis [10,14–21]. Doing so may result in single source DNA profiles for all contributors as well as new ‘mini mixtures’ with less complexity compared to the original bulk mixture due to a decrease in the NOC [10]. In this study, this concept was applied to complex mixture analysis using the previously reported direct single cell subsampling (DSCS) method, [10] but now, analyzed with PG methods validated for enhanced DNA-STR typing of single (or few) cells.

## 2. Methods

### 2.1. Sample collection

Buccal swabs were collected from volunteers by swabbing the inside of the mouth and cheek with a sterile cotton swab according to procedures approved by the University’s Institutional Review Board.

### 2.2. Slide creation

#### 2.2.1. Gel-Film slide creation

Gel Film microscope slides were created by attaching Gel-Pak^®^ Gel-Film^®^ (WF, x8 retention level) (Hayward, CA, USA) to clean glass microscope slides by the Gel-Film adhesive backing. The clear protective covering was removed when creating mixture slides.

#### 2.2.2. Mixture slide creation

Freshly collected or dried/frozen buccal swabs from individual donors were agitated in 300 μL of TE^-4^ buffer. The resulting solutions were centrifuged at 300 RCF to create an epithelial cell pellet. The supernatants were discarded, and 300 μL of TE^-4^ buffer was added to resuspend each cell pellet. The Countess™ II FL (ThermoFisher Scientific) automated cell counter was then used according to manufacturer recommendations to determine the cell concentration of each of these cell suspensions. Equals concentrations of each donor cell suspension were mixed to create the desired mixtures (ie 2-, 3-, 4-, 5-, or 6-person). Mixture cell suspensions were stored frozen. Once the desired mixture was created, 60 μL of the mixture solution was pipetted onto the Gel-Film^®^ microscope slide and spread out using a sterile swab. The slide was then stained with Trypan Blue for 1-2 min and gently rinsed with nuclease-free water. Slides were air-dried overnight.

#### 2.2.3. 3M^TM^ adhesive slide creation

Prior to cell collection, a 3M^TM^ (St. Paul, MN, USA) adhesive slide reservoir was made. Double-sided tape was used to attach the 3M^TM^ adhesive to a clean glass microscope slide, and the adhesive backing was removed. The slide was then stored in a desiccator until needed [22,23].

### 2.3. Cell recovery

Cells were visualized (from the gel-film mixture slide previously created in section 2.2.2) using a Leica M205C stereomicroscope (190-240X magnification) and collected directly into sterile 0.2 mL PCR flat-cap tubes containing Prep-n-Go^TM^ Lysis buffer (ThermoFisher Scientific, Carlsbad, CA, USA). This was done via a sterile tungsten needle and water-soluble 3M^TM^ adhesive. The tungsten needle was used to obtain a small ball of 3M^TM^ adhesive (from the adhesive slide reservoir created in section 2.2.3) which was then used to adhere selected visualized cells from the mixture slides [22,23]. Still under the microscope, the adhesive-tipped tungsten needle containing adhered cells was inserted into an amplification tube (0.2 mL) containing 1 μL lysis buffer until the 3M^TM^ adhesive was observed to solubilize. Twenty 1- and 2-cell subsamples were first collected and analyzed for every mixture. If the expected minimum number of contributors (as determined from bulk analysis), or if probative log (LR) values were not obtained for all donors, additional s=10 x 1 and 2 cell subsamples were collected until probative results were achieved. Twenty 1 and 2-cell samples were collected for the 1:1 and 1:1:1 mixtures. Thirty 1 and 2-cell subsamples were collected for the 1:1:1:1 and 1:1:1:1:1:1 mixtures and forty 1 and 2-cell subsamples were collected for the 1:1:1:1:1 mixture.

### 2.4. Direct lysis/ autosomal short tandem repeat (STR) amplification of cells

Cells were collected directly into 1 μL Prep-n-Go^TM^ lysis buffer as described in section 2.3. Samples were then incubated at 90°C→ 20 mins; 25°C→ 15 mins. After lysis of cells, the samples were amplified using the GlobalFiler™ Express kit (ThermoFisher Scientific, Carlsbad, CA, USA) with a reduced reaction volume of 5 μL and increased cycle number (32 cycles). The GlobalFiler™ Express reaction mix was prepared consisting of 2 μL PCR mix and 2 μL primer mix added to the 0.2 mL PCR flat-cap tubes. Samples were amplified using a protocol of 95°C→ 1 min; 32 cycles: 94°C→ 3 sec, 60°C→ 30 sec; 60°C→ 8 mins; 4°C→ hold. Positive (1 μL 31.25 pg 007) and negative amplification controls (0-cell samples and amplification blanks) were included in each amplification batch.

### 2.5. Donor reference samples and bulk mixtures

#### 2.5.1. DNA isolation and quantitation

DNA was extracted from reference buccal swabs and 60 μL of each mixture cell suspension using the QIAamp DNA Investigator kit (QIAGEN, Germantown, MD, USA) according to the manufacturer’s manual extraction procedure. An extraction blank was included in each extraction set. Quantifiler^®^ Duo DNA Quantification kit (ThermoFisher Scientific) was used to quantify extracts according to the manufacturer’s recommended protocols using the Applied Biosystems’ 7500 real-time PCR instrument (ThermoFisher Scientific).

#### 2.5.2. Autosomal STR amplification (reference samples and bulk mixtures)

Amplification of reference samples and bulk mixtures were conducted using the GlobalFiler™ kit according to manufacturer recommended protocols. One nanogram of input DNA was used. The amplification protocol was: 95°C→ 1 min; 29 cycles: 94°C→ 10 sec, 59°C→ 90 sec; 60°C→ 10 mins; 4°C→ hold. A positive and negative amplification control was included in each amplification.

### 2.6. PCR product detection

One microliter of GlobalFiler™ or GlobalFiler™ Express amplified product was added to 9.5 μL Hi-Di™ formamide (ThermoFisher Scientific) and 0.5 μL GeneScan™ 600 LIZ^®^ size standard (ThermoFisher Scientific). Samples were injected on an Applied Biosystems’ 3500 Genetic Analyzer using POP-4^TM^ polymer and Module G6 (15 s injection, 1.2 kV, 60 °C). Samples were analyzed using GeneMapper v1.6 software (ThermoFisher Scientific).

### 2.7. Probabilistic Genotyping (PG)

#### 2.7.1. Standard Bulk Mixture Probabilistic Genotyping

Probabilistic Genotyping Software STRmix™ v2.8 was validated for use with standard bulk mixtures and reference samples. Each bulk mixture was analyzed according to the known number of donors. For example, for a 3-person mixture, the likelihood ratio was conditioned as known donor 1 and 2 unknown individuals contributing to the mixture vs. 3 random unknown individuals. This approach was repeated for the remaining two donors. Probabilistic Genotyping was also used to analyze all reference profiles as single source to determine the random match probability. The sub-source log (LR) was reported. The five and six-person bulk mixtures were run with the assistance of STRmix™ Technical and Scientific Support (Environmental Science and Research) and a beta STRmix™ v2.9 for the 6-person mixture specifically. The FBI Caucasian database was used for all allele frequencies in all mixture experiments.

#### 2.7.2. DSCS Probabilistic Genotyping

Probabilistic Genotyping Software STRmix^TM^ v2.8 was validated for use with 1 or 2-cell subsamples. Each 1-cell subsample was run as a single source sample and the log(LR) reported. Two-cell subsamples were either run as single source samples or, when appropriate, 2-person mixtures. A database of N = 1,000 known non-contributors provided by STRmix^TM^ support (from their training courses) was also utilized. Additional in-house analyst genotypes were added to the database to monitor for contamination.

##### 2.7.2.1 DSCS Analytical Threshold

The analytical threshold used when analyzing cell subsamples collected by DSCS was calculated by analyzing 30 negative controls (0-cell collections and amplification blanks) and averaging the highest peaks seen at each dye channel. The equation AT= 2*(highest peak – lowest through) was used [24]. (Blue = 53 RFU; Green = 86 RFU; Yellow = 46 RFU; Red = 63 RFU, and Purple = 58 RFU)

##### 2.7.2.2 DSCS Drop-in Rate

The drop-in rate was calculated by analyzing 35 negative controls for the presence of drop-in alleles. The equation used was the number of drop-in events/ (the number of loci scored x the number of samples). The drop-in data did not fit a gamma distribution as limited drop-in data was available. Therefore, a uniform distribution was utilized (graph not shown). A drop-in rate of 0.0164 and a drop-in cap of 30,000 RFUs was used to allow for any height allele to be considered as possible drop-in. (This is an increase compared to the parameters used for the standard bulk mixtures (bulk mixture: drop-in rate= 0.0001 & cap= 100 RFU)).

##### 2.7.2.3 DSCS STRmix™ Stutter Parameters

Stutter regression files were created for each locus and each type of stutter encountered according to the STRmix™ implementation and validation guide. The STRmix^TM^ stutter parameters were then determined using the Model Maker function. Model Maker utilized 161 x −1, −2, −3, −4, or 5-cell subsamples to model the stutter distribution. The maximum allowed stutter percentages were larger than those typically seen with standard analysis because of elevated stutter that is characteristic of low template stochasticity. The DSCS stutter parameter results are shown in Supplementary Table 1. For comparison, the stutter parameters used for the standard bulk mixtures are provided in Supplementary Table 2.

##### 2.7.2.4 DSCS STRmix™ Post Burn-in Accepts

A single source 3-cell subsample was analyzed 5 times using 5,000, 50,000, and 500,000 post burn-in accepts. It was determined that 500,000 post burn-in accepts gave the most consistent log (LR) results (graph not shown) for DSCS samples. Additional DSCS settings (including, for comparison, those for standard bulk mixtures) are provided in Supplementary Table 3.

#### 2.7.3. DSCS Probabilistic Genotyping Replicate Analysis

After PG analysis of each subsample, replicate analysis [25] was performed per donor for single source 1-cell samples, single source 2-cell samples, and 2-cell mini-mixtures. The top up to six 1-cell subsamples that gave a log (LR)s ≥ 1 per donor were utilized for 1-cell replicate analysis. These 1-cell subsample replicates were run as single source ie. LR= Donor 1 vs. unknown individual. Likewise, the top up to six single source 2-cell subsamples that gave log (LR)s ≥ 1 for a donor were utilized for 2-cell replicate analysis. Two-cell mini-mixture replicates were conducted any time a log (LR) ≥ 1 was achieved for a donor regardless of identity of the second donor.

### 2.8 Description of DSCS Method

An infographic of the DSCS method for complex mixture analysis that uses an equimolar 4-person mock mixture for illustrative purposes is provided in Fig. 1. The standard bulk mixture analysis, which begins with the extraction and *de facto* homogenization of DNA from a mixed stain, results in an admixture of DNA from contributors where each contributor’s genotype cannot be distinguished (Fig. 1, left side). Direct single cell subsampling of the same mixture allows for collection of multiple 1-2 cell subsamples. This allows for single source profiles of all individuals (A, B, C, and D) from the mixture to be obtained by 1-cell subsamples as well as some 2-cell subsamples. By increasing the number of cells collected in subsampling from 1 to 2, the amount of input DNA doubles thus increasing the probability of achieving a full profile if both cells originate from the same donor. However, some 2-cell subsamples still result in admixed profiles (i.e. 2-cell mini-mixtures) which are less complex than the original bulk mixture due to a decrease in the number of contributors (Fig. 1, right side).

**Figure 1.**
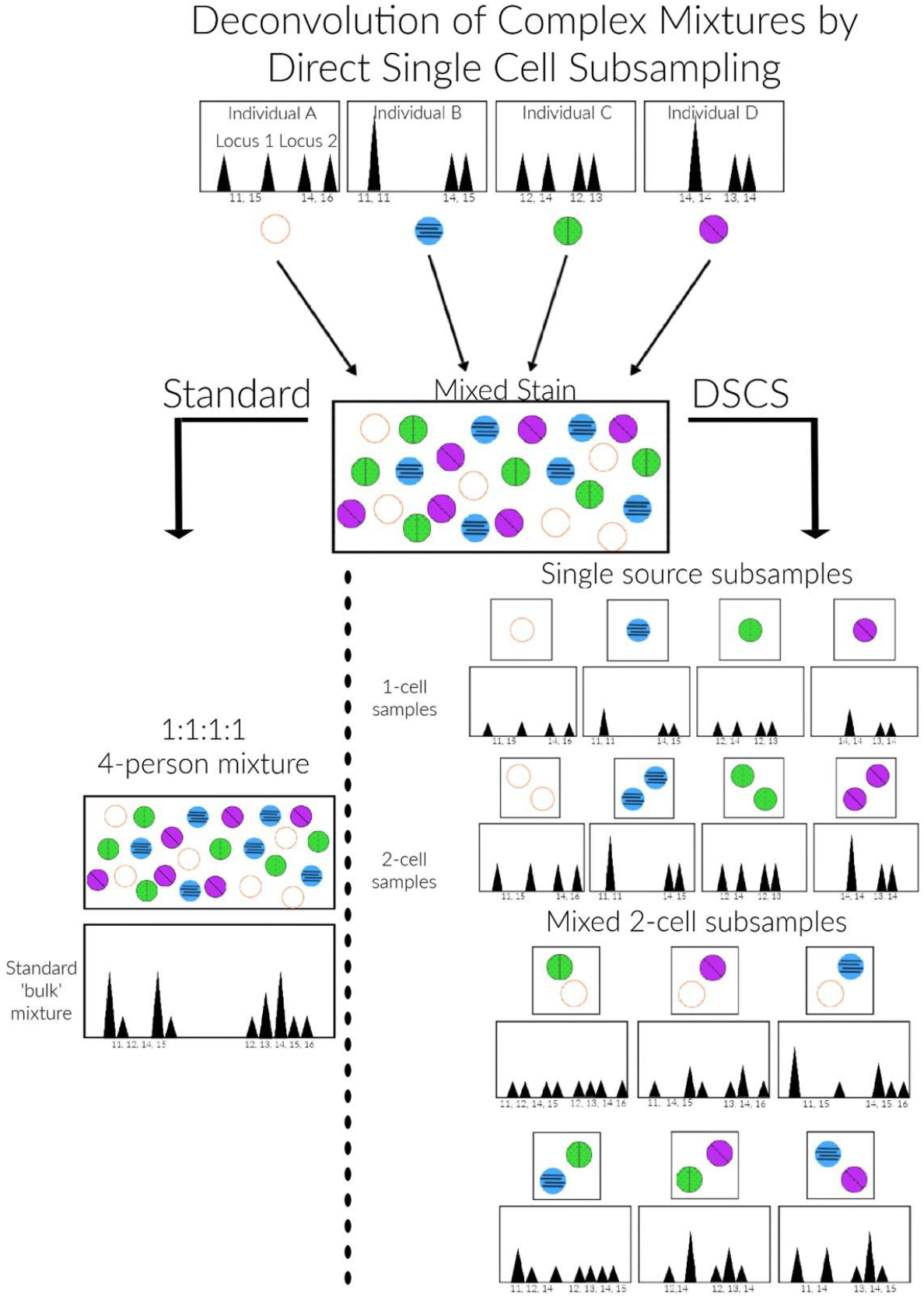
Direct Single Cell Subsampling (DSCS) scheme for complex mixture analysis.

The direct single cell subsampling method produces a number of different subsample types that are then analyzed with probabilistic genotyping (PG). Fig. 2 shows an infographic summarizing the PG analysis decision tree developed in this study for the subsample types using the same 4-person mock DNA mixture shown in Fig. 1. Prior to DSCS analysis, a bulk mixture is analyzed using standard methods. With DSCS, a collection of single cell subsamples results in single source profiles of donors to the mixture. These individual subsample DNA profiles from single cells are then analyzed using PG techniques validated for use with direct PCR, reduced reaction volumes and few cells. For this, each recovered subsample genotype is PG tested individually against all potential donors comprising the mixture. Thus, for each subsample a number of LRs, dependent upon the number of potential donors tested as a contributor, will be calculated. Any subsample profile that gives a log (LR) ≥ 1 for a specific donor is then deemed suitable for PG replicate analysis for that specific donor, and the presence of at least one other subsample with a positive log (LR) for the same donor is needed for PG replicate analysis. By this process the genotyped subsamples from the mixture are essentially categorized according to putative donor prior to conducting the replicate analysis. The same process is conducted for 2cell single source subsamples. For 2-cell mixture samples, i.e. ‘mini-mixtures’, replicate analysis is conducted when multiple mixture samples give a log (LR) ? 1 for a specific donor. For simplicity, the subsequent use of the term DSCS in this report is taken to mean the PG validated DSCS method.

**Figure 2.**
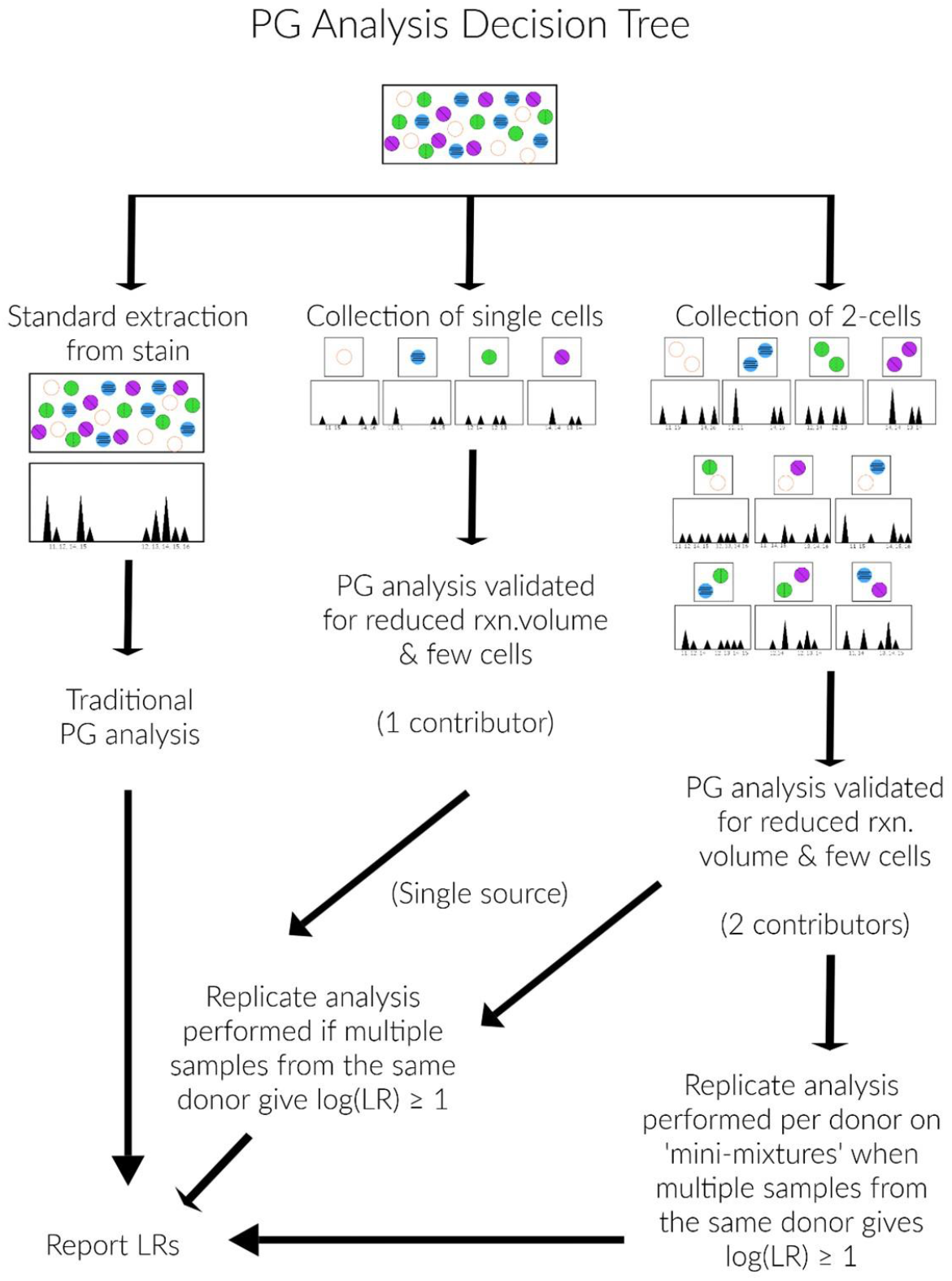
Subsample types and associated probabilistic genotyping (PG) analysis method decision tree.

## 3. Results

### 3.1 STR kits/lysis conditions

For DSCS, more discriminatory and sensitive commercial STR kits than previously reported [10] were tested using 1-, 2-, 3-, 4-, 5- and 10-cell subsamples. Based upon the quality of the profiles and the number of alleles recovered (Supplementary Fig. 1), two kits were found to perform particularly well, namely GlobalFiler^®^ Express (GFE) and PowerPlex^®^ Fusion 6C (PPF6C). GFE was chosen as the system of choice for PG validation in this study since it performed best for single buccal cells and was designed specifically for direct PCR, which is used in the DSCS process. However, PPF6C represents a suitable alternative sensitive STR system that can be used for single cell STR typing. Examples of GlobalFiler^®^ Express 1 and 2 cell EPGs can be seen in Supplementary Figs. 2, 3, and 4.

### 3.2 PG validation

The PG software system STRmix™ was validated using GlobalFiler^®^ (GF) for standard mixture analysis according to the recommendations of the software developers [5,26]. For use with DSCS, PG validation was conducted using the same methodology except with subsamples of discrete nucleated cells as input (1-5 cells, equivalent to approximately 6.6 pg-33 pg DNA) typed with GlobalFiler^®^ Express (GFE) in contrast to standard applications which utilized DNA input (~ 0.1-1.0 ng gDNA) arising from dilutions of bulk DNA extracts. The STRmix™ validation comprised two phases, namely the determination of empirically derived parameters used to formulate a model and subsequent testing of the sensitivity and specificity of the model to deconvolute known mixtures.

As an additional test of the accuracy of the PG parameters for 1-5 cells, a 3-cell single source sample was analyzed using probabilistic genotyping, and then, an artificially degraded profile (created by decreasing the peak heights of the high molecular weight alleles by 80% and the low molecular weight alleles by 5%), an artificially inhibited profile (created by decreasing the peak heights of D22S1045, D21S11, D13S317, and D2S1338 loci by 40%), and a profile with artificial drop-in (created by adding a drop-in 8 allele at D5S818 with 15,000 RFUs) were created and analyzed. A single dye channel from the degraded, inhibited, and drop-in profiles is shown in Supplementary Fig. 5. In all 3 artificial profiles, PG reported the same log (LR) as the original sample (Supplemetary Table 4).

### 3.3 Testing the limits of standard PG analysis with complex mixtures

With the PG systems validated for standard DNA mixture analysis (i.e. ‘bulk’ analysis rather than subsampling by DSCS) we tested the limits of a commonly used PG system in extracting probative information from standard mixtures. To that end we selected random 2, 3, and 4 person DNA mixtures (50 from each mixture class) from in-house prepared DNA mixtures and the PROVEDIt database [27,28] and calculated the log (LR)s for each known donor using STRmix™ The mixtures comprised a variety of donors and mixture weight ratios. Quantitatively, in these 2-,3-, and 4-person mixtures, only 86%, 74%, and 32% of each respective mixture type gave very strong support (log(LR) ≥6) for each contributor (Table 1).

**Table 1.** Percentage of standard bulk mixtures in which all donors reach the specified threshold.

| Mixture Type (n=50) | $\log(LR) > 0$ | $\log(LR) \geq 6$ |
| --- | --- | --- |
| 2-person | 94% | 86% |
| 3-person | 98% | 74% |
| 4-person | 88% | 32% |

Note that a similar number of 5- and 6-person mixtures, albeit 5-person mixtures are available on the PROVEDIt database, could not return LRs in this study due to computational speed and memory limitations. Indeed, we were able, with the help of a supercomputer and STRmix™ Technical and Scientific Support (Environmental Science and Research), to perform standard PG analysis on an in-house generated 5- and 6-person mixture. In both mixtures, the LRs from three of the known contributors failed to reach the log (LR) = 6 threshold.

These results indicate that the DSCS approach, if successful, could be of benefit in recovering more highly probative genotype information from each of the contributors to complex mixtures.

### 3.4 Sensitivity and Specificity of DSCS

Since DSCS is dependent upon analysis of low template samples, a high degree of dropout is observed across the 21 GFE STR loci (Table 2). However, 1-cell subsamples recovered on average 18 alleles while 2-cell single source subsamples recovered 26. This high degree of dropout can occasionally result in profiles with high allele counts but low LRs principally due to single allele drop-outs from heterozygous loci. Since the simultaneous deconvolution of replicates within a single analysis reduces the impact of stochastic effects [25], the available replicate analysis capability with PG systems was further investigated for DSCS applicability.

**Table 2.** Percentage of Allele Drop-out per Locus.

| Allele Drop-out |  |  |
| --- | --- | --- |
| Locus | 1 Cells | 2 Cells |
| D3S1358 | 52% | 25% |
| vWA | 59% | 39% |
| D16S539 | 68% | 44% |
| CSF1PO | 66% | 45% |
| TPOX | 74% | 54% |
| AMEL | 36% | 24% |
| D8S1179 | 50% | 31% |
| D21S11 | 67% | 38% |
| D18S51 | 61% | 39% |
| D2S441 | 36% | 18% |
| D19S433 | 39% | 25% |
| TH01 | 57% | 34% |
| FGA | 55% | 35% |
| D22S1045 | 40% | 20% |
| D5S818 | 48% | 35% |
| D13S317 | 66% | 41% |
| D7S820 | 67% | 46% |
| SE33 | 75% | 56% |
| D10S1248 | 43% | 25% |
| D1S1656 | 52% | 36% |
| D12S391 | 66% | 51% |
| D2S1338 | 75% | 51% |
| Total | 58% | 38% |
| Avg. Alleles | 18 | 26 |

The sensitivity of PG for single cell genotyping by DSCS was assessed. Sensitivity is defined in PG analysis as the proportion of true contributors that return positive LR values (i.e. lends support for the inclusionary proposition H_1_ or H_p_). The LR threshold for most sensitivity studies has traditionally been set at 1 (i.e. log (LR) =0) and thus any sample that returns a log (LR) > 0 would be regarded as a positive for the sensitivity analysis. In this work, since the aim is to better recover highly probative donor genotype information from complex mixtures, we also assessed the ability of the DSCS procedure to achieve a minimum log (LR) of 6 for individual cell subsamples from each mixture contributor. We chose this latter LR threshold since it excludes the possibility of obtaining false positive results from (unrelated) non-contributors, and represents the upper ‘evidentiary significance’ threshold beyond which the international forensic community has decided no additional verbal expressions beyond providing “extremely strong support” for the inclusionary proposition are applied [29,30]. Equimolar 2-6 person mixtures were prepared from buccal cells as described (Methods Section 2.2.2) and a total of 291 cell subsamples (110 x 1 cell subsamples, 81 x 2-cell single source subsamples, 100 x 2-cell mini mixtures) were recovered by DSCS and subjected to PG analysis with STRmix. Table 3 shows the determined DSCS specificity obtained using two thresholds, namely the traditional log (LR) > 0 and the log (LR) > 6 targeted in this work.

**Table 3.** Percentage of subsamples and replicates analyzed with STRmix™ that reach the specified threshold.

| Sample Type | Subsample<br>log(LR)>0 | Replicate | Subsample<br>log(LR)≥6 | Replicate |
| --- | --- | --- | --- | --- |
| SS 1 cell (s=100) | 68% | 100% | 26% | 94% |
| SS 2 cell (s=81) | 84% | 100% | 60% | 93% |
| Mix 2 cell (s=110) | 60% | 100% | 29% | 86% |

The traditionally measured specificity threshold (log (LR) > 0 threshold) was met by the majority of 2-cell single source subsamples (84%), 1-cell subsamples (68%), and 2-cell mini mixture subsamples (60%). While some of the subsamples were not of a high enough quality to reach the ‘very strong support’ threshold for the inclusionary hypothesis (log (LR) > 6), the majority (60%) of 2-cell single source subsamples, 26% of 1-cell subsamples, and 29% of 2-cell mini mixture subsamples did so. Significantly, some 1 and 2-cell single source subsamples returned log (LR)s ≥ 21 (3% and 19% respectively), which is close to the inverse of the match probability for the donors. The latter effectively represents the upper bound of the achievable LR [31] and therefore also the maximum information recoverable from the subsample.

Replicate analysis was performed in the hope and expectation that it might improve the sensitivity of DSCS analysis. Cell subsamples are low template DNA samples and replicate analysis of low template samples has been shown to not only reduce characteristic confounding stochastic effects [32] but to be able to obtain higher LRs from multiple low template replicates than from a single bulk sample [31]. Sensitivity was significantly improved (i.e. LRs increased) when replicate analysis was performed on the DSCS subsamples. Table 3 shows the results when the same samples as above were subjected to replicate analysis for which, on average, 4 subsamples (± 1) exhibiting the highest returned LRs were used. Replicate analysis resulted in 100% sensitivity for all subsample types when the threshold was set at log (LR) > 0. In almost all instances, returned LRs provided ‘very strong support’ (i.e. log (LR) > 6) for the inclusionary hypothesis (94 % for 1-cell replicates, 93% for 2-cell single source replicates, and 86% of 2-cell mini mixture replicates). Significant proportions of 1- and 2-cell single source replicates resulted in log (LR)s ≥ 21 (41% for 1-cell replicates, 79% for 2-cell replicates) (data not shown).

Specificity is the proportion of true non-contributors that return LR < 1 values (i.e. log (LR) < 0) and thus supports the exclusionary proposition (H2 or Hd). The specificity of DSCS analysis was checked for each single source 1-cell and 2-cell subsamples used in the sensitivity analysis. This was done by substituting 1000 random known non-contributor DNA profiles instead of the known contributors (i.e. substituting the non-contributor for the known in the inclusionary proposition (i.e. H_1_ or H_p_)) and calculating the LR for each. The results are shown in Figure 3 in which the log (LR) values for known donors are plotted for each subsample with respect to the number of alleles detected. Thus, each single source 1 and 2 cell subsample (blue and yellow respectively) will return a single LR for the known contributor and 1000 LRs for the non-contributors (orange). Likelihood ratios of 0 were plotted as a log (LR) = −50. As before, the log (LR) achieved from known donors to single source subsamples increased as allele recovery increased while the number of non-contributor false positives with an LR > 1 also decreased as allele count increased. As can be seen from Fig. 3, non-contributors never exceeded the 10^6^ LR threshold (dashed line) although several cells with less than 10 alleles returned LRs >1. The majority of false positives had a log (LR) between 1 to 3 indicating ‘uninformative’ or ‘limited support’ [29,30] but one non-contributor subsample did result in a log (LR) of 5. There was good separation between the log (LR)s obtained from known and non-contributors when the detected number of alleles from the cell(s) exceeded 15. Similarly, the log (LR)s achieved with known donors and known non-contributors to 2-cell mini mixture subsamples were plotted according to the number of non-shared alleles. No false positives were obtained at or above a threshold of log (LR) = 6 (Supplementary Fig. 6).

**Figure 3.**
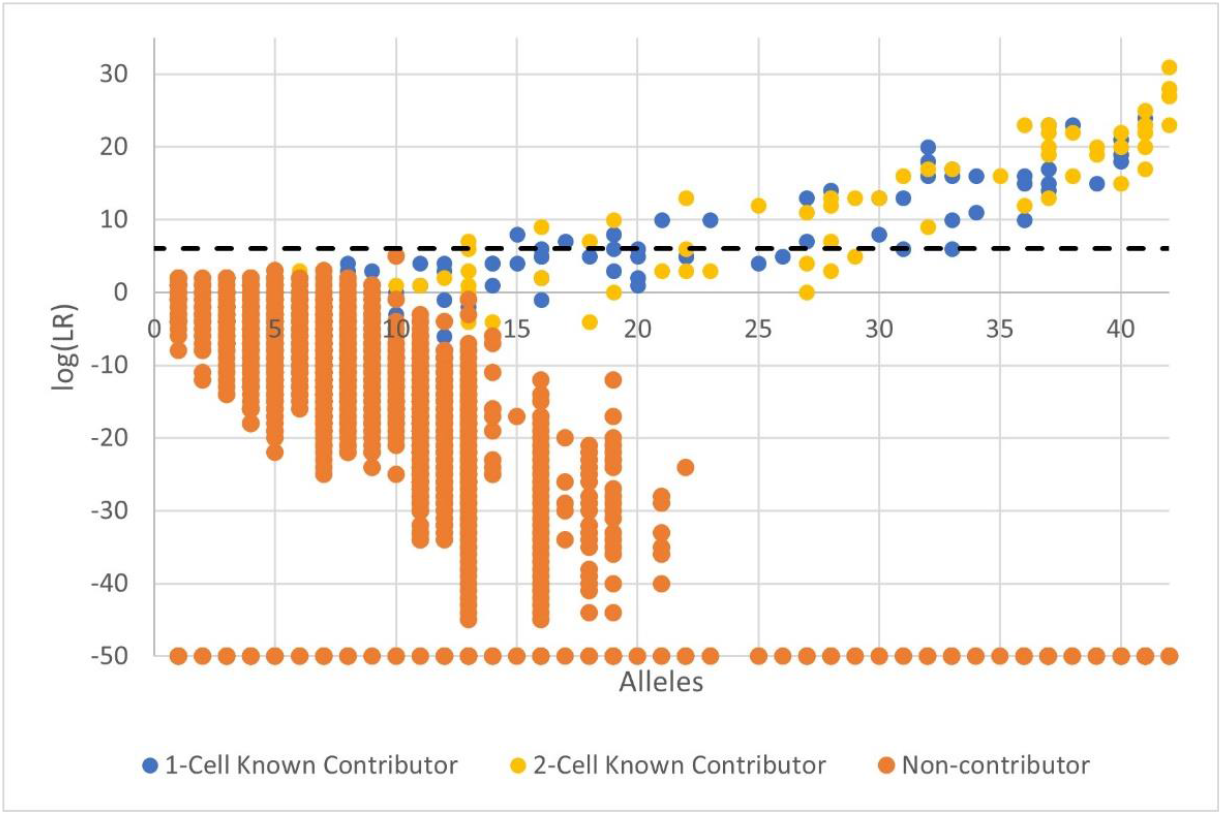
Sensitivity and specificity of DSCS (STRmix™) analysis. Single source subsamples. LR = 0 plotted as −50. Blue circles (1-cell known contributor), yellow circles (2-cell known contributor), orange circles (non-contributor)

Improved specificity and sensitivity of DSCS was obtained by replicate analysis (Fig. 4). A plot of log (LR) from the original single source cell subsamples versus log (LR) replicates indicated that, in almost every instance, replicates increased the recovered LR from known contributors (i.e. sample coordinates located above the diagonal). Simultaneously the false positive LR rate was reduced to zero (i.e. no non-contributor samples now returned an LR>0). Similarly, replicate analysis of the 2-cell mini-mixtures resulted in improved LRs compared to individual subsamples (Supplementary Fig. 7). The highest false positive LR seen with mini-mixture replicates was a log (LR) = 1.7 which would be classified as uninformative according to the SWGDAM recommendations for verbal qualifiers.

**Figure 4.**
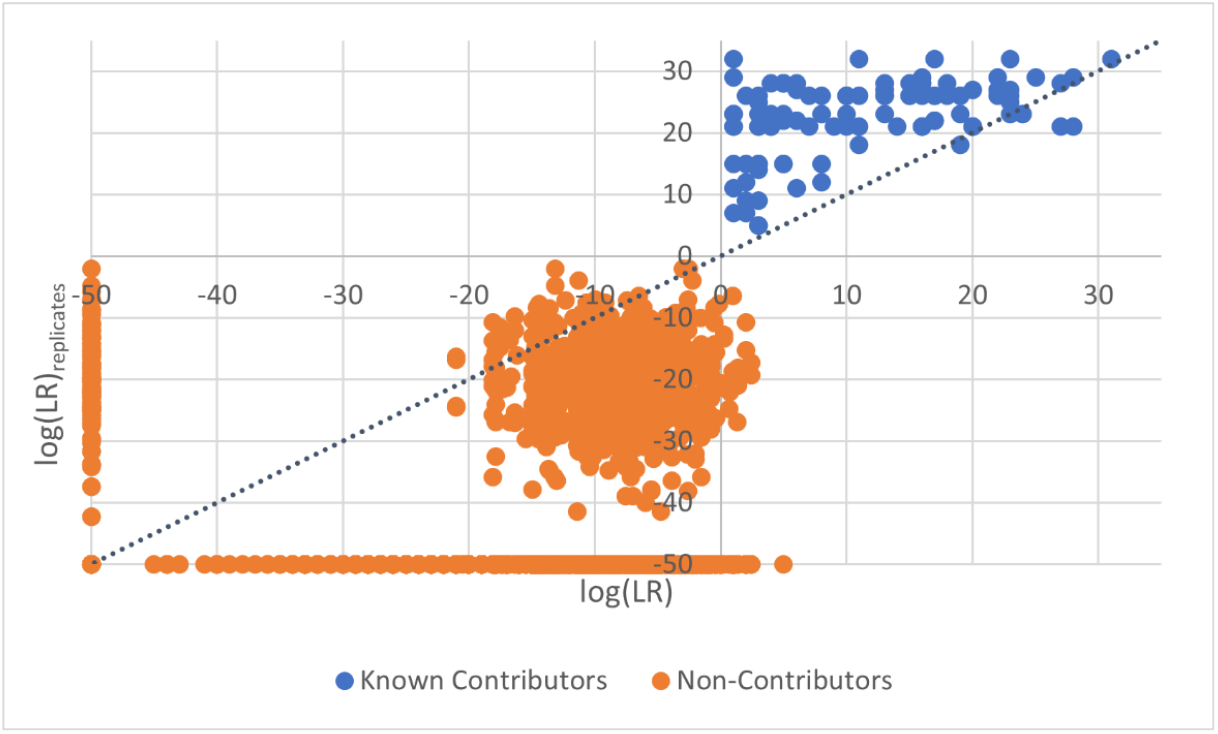
Specificity improvement of DSCS (STRmix™) with replicate analysis. Known contributors above the diagonal (i.e. y=x) represent improved LR recovery from replicate analysis. Single Source Cell Replicates. LR = 0 plotted as −50. Blue circles (known contributors); orange circles (non-contributors)

In summary the above results indicate that the specificity and sensitivity of DSCS analysis increases with replicate analysis and can result in the recovery of most (sometimes all) donor genotype information.

### 3.6 DSCS applied to complex mixtures

The extent to which DSCS minimized genotype information loss from each of the donors to complex mixtures was investigated using equimolar, buccal 2-6 person admixtures. The replicate log (LR)s from DSCS analysis was determined as previously described (section 2.7.3) for each contributor by using up to 6 log (LR) ≥1 subsamples (i.e. the highest six log (LR)s or, if less than 6 replicates with a log (LR) > 1 are obtained, all replicates are used). Supplementary Figs. 8 and 9 shows vertical bar graphs of the replicate analysis log (LR) results for each contributor per mixture (heights of the bars) as well as the individual subsample log (LR) results that were used as replicates (dots within contributor bars). Across all mixtures shown the average number of subsample replicates used in the replicate analysis per donor per mixture was 4, with a median of 3.

The DSCS replicate log (LR)s were compared to those from the ground truth single source samples (i.e. log (1/RMP from reference samples)) and that obtained from standard bulk mixture DNA analysis using STRmix^TM^ (Figs. 5 and 6). DSCS recovered more genotype information from every donor in every mixture compared to standard PG mixture analysis, with the LR gain being noticeably more significant in the 5 and 6 person mixtures. One of the donors to the 4-person mixture (FM20), three donors to the 5-person mixture (S3, S5 and FM20), and three donors to the 6-person mixture (S8, CM31 and SA10) failed to return a log (LR) ≥6 with standard analysis whereas DSCS returned highly probative log (LR)s from the same donors (range 8-27). In some donors, recovery of their full intrinsic genotype information (i.e. log (LR)DSCS = log (1/RMP)) was achieved. An additional 6-person mixture was analyzed as can be seen in Supplementary Fig. 10. and highly probative log (LR)s were recovered by DSCS from every donor (range 13-24). No standard bulk mixture analysis results were available for comparison with this mixture due to computational limitations.

**Figure 5.**
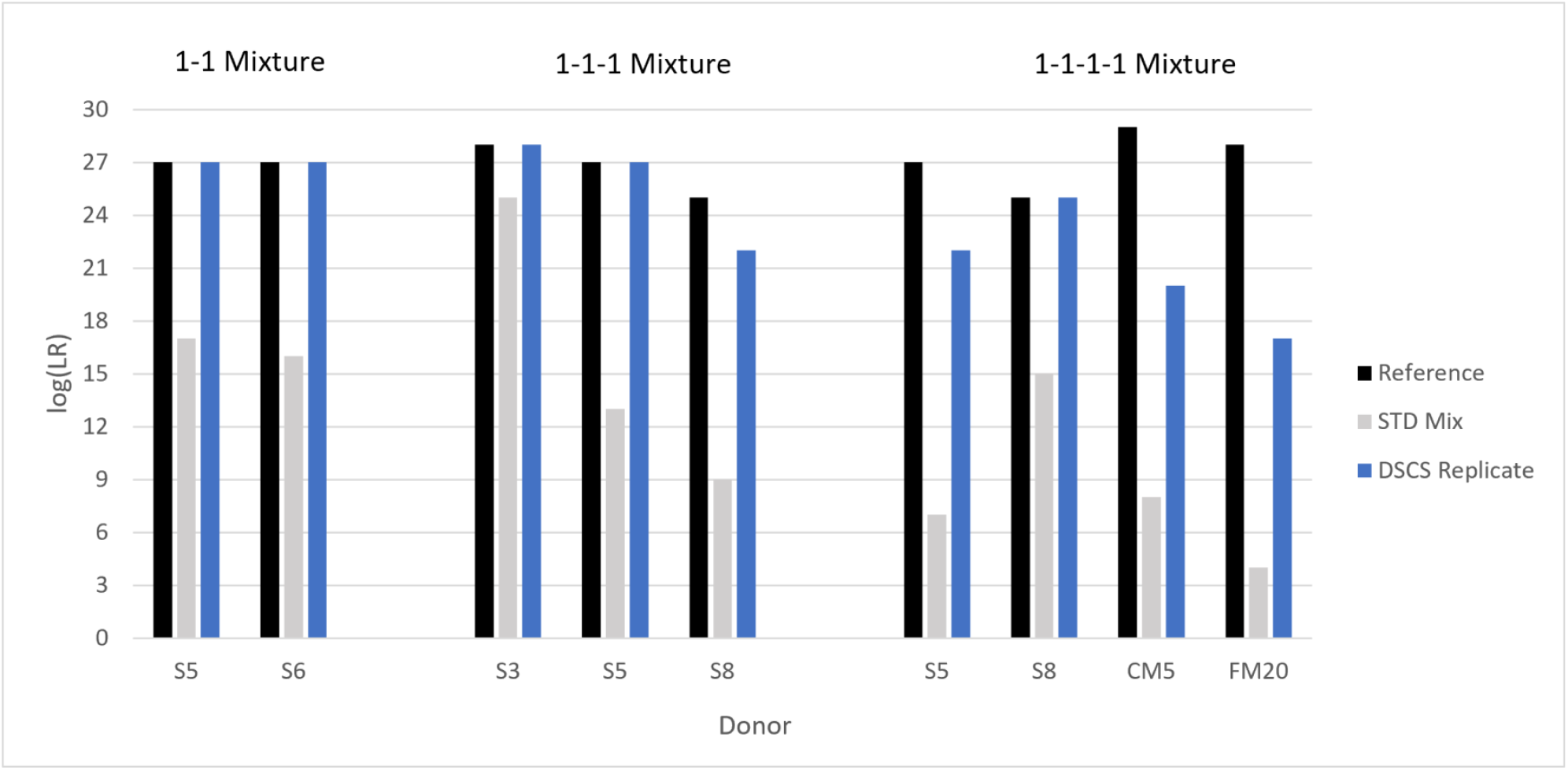
Increased contributor log(LR) recovery in 2-4 person mixtures by DSCS (STRmix™) compared to standard PG mixture analysis.

**Figure 6.**
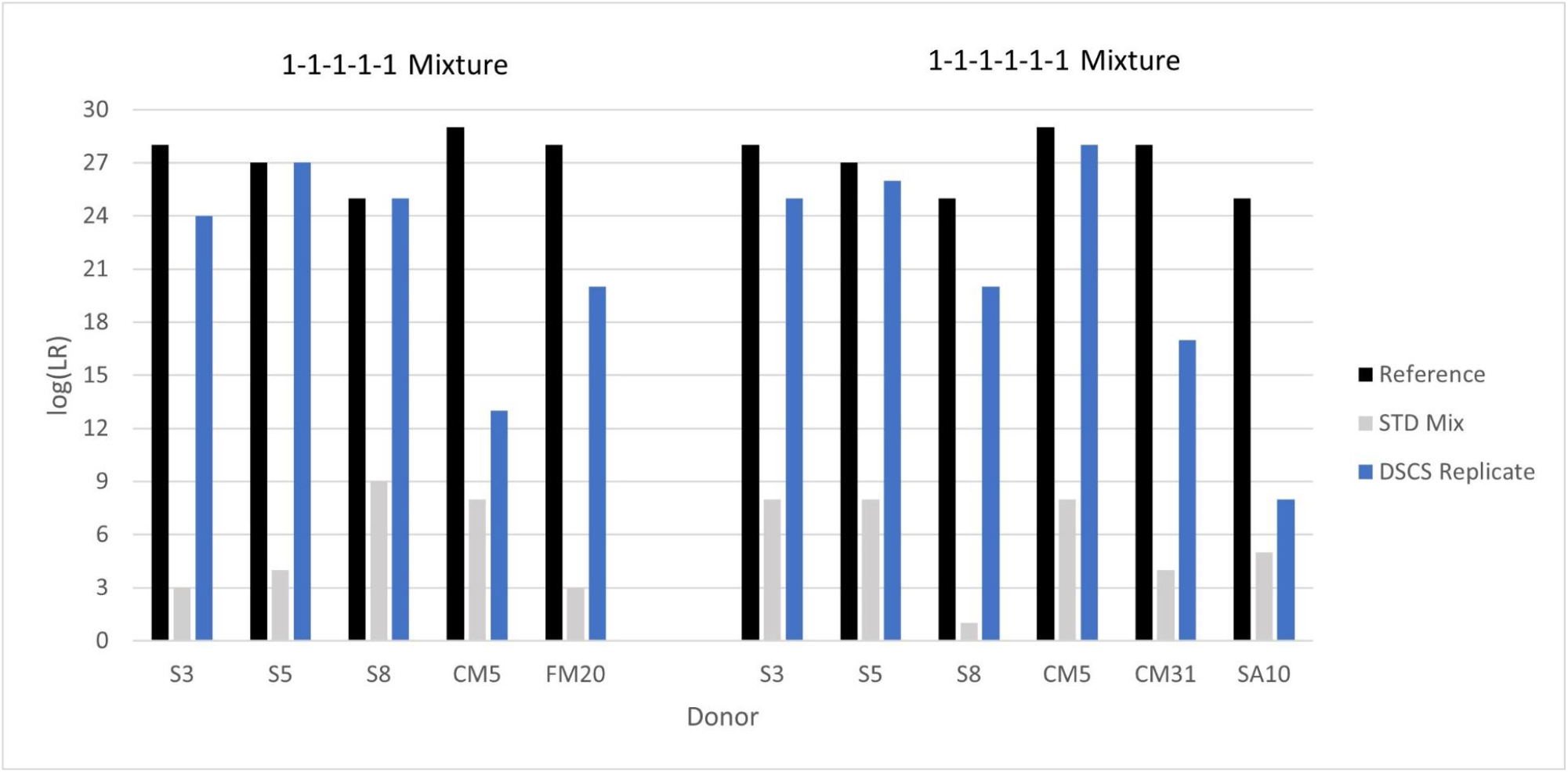
Increased contributor log(LR) recovery in 5-6 person mixtures by DSCS (STRmix™) compared to standard PG mixture analysis.

## Discussion

We reasoned that the use of continuous probabilistic genotyping (PG) methods and models, which use empirically derived variance parameters and can perform replicate analysis, could be productively employed in single cell analysis to provide accurate and reliable genotype resolution and LR (i.e. evidential weight) estimates. In this proof-of-concept (PoC) work, we have shown that direct single cell subsampling (DSCS) of complex DNA mixtures, combined with high sensitivity DNA typing followed by PG interpretation, can recover more genotype information (as demonstrated by increased LRs) from each of the contributors than that obtained by standard methods. We define complex mixtures in this study as samples that contain DNA from two or more contributors in which stochastic effects or allele sharing cause uncertainty in determining contributor genotypes [12]. The information gain was achieved because single-cell subsamples from the mixtures resulted in deconvoluted single-source DNA profiles while 2-cell subsamples returned either single-source DNA profiles or new mini-mixtures that are less complex than the original mixture due to the presence of fewer contributors. A suitably validated PG software system, STRmix™, was implemented to analyze the original bulk mixtures, singlesource cell subsamples, and the 2-cell mini mixture subsamples obtained from the 2-6 person complex mixtures studied. In every mixture, the DSCS approach gave improved results for each donor compared to standard bulk mixture analysis. With 5- and 6-person complex mixtures, DSCS recovered highly probative LRs (> 10^20^) from donors that had returned non-probative LRs (<10^3^) by standard methods. In many such instances, the returned donor likelihood ratios (LRs) were comparable to that seen in their single-source reference profiles. We employed a different single cell-validated PG system, EuroForMix (v3.1.0, Quantitative LR MLE based) [9,33] to analyze the mixtures. The two different PG software systems returned remarkably similar individual contributor LR values with the DSCS samples with no significant discordant results (data not shown). This represented an additional diagnostic check [34] on the accuracy and reliability of the DSCS method’s performance with the sample set studied. The DSCS method with replicates represents a better and more objective approach to donor assignment and interpretation of single cell subsamples than previous methods [10] which principally used manual consensus profiling from replicates to recover genotype information.

Maximum information gain was achieved by taking advantage of replicate contributor subsamples that are intentionally obtained by the DSCS process and the subsequent use of the replicate analysis capability of the PG systems. Consensus DNA profiles obtained from low template analysis of bulk extracted samples, given enough replicates, can reconstitute much of the genotype information that would otherwise be lost due to low template DNA stochastic processes [32,35]. Here we employed replicate analysis to do the same thing for DSCS single cell subsamples using the replicate analysis capabilities of the PG systems. The way replicates work here is that cell subsamples are genotyped and clustered/classified essentially according to genotype, by performing PG analysis on all individual subsamples that yield STR profiles and determining which, if any, return an LR ≥ 1 for any of the potential POIs (Fig. 2). In order to test the PG system’s replicate accuracy, whereby a subsample might be misclassified for replicate use, simulations were carried out in which a single subsample (with LR <1) collected from the same bulk mixture but from a different donor (i.e. incorrect donor for the replicates being analyzed) was added to a replicate subset originating from a particular donor and replicate analysis performed: this was repeated by substituting other single subsamples with an LR < 1 (data not shown). When including misclassified samples for single source replicate analysis, if a poor-quality profile was used (~4 alleles from a possible 42), relatively no change was seen in the log (LR) achieved. As profile quality of the misclassified subsamples increased, positive log (LR)s of the replicates were no longer obtained (LRs of 0 or strongly negative). When including misclassified subsamples for mini-mixture replicate analysis, sometimes, a positive result for the true donors was still seen, but it did not increase the LRs obtained without the use of replicate analysis. Thus, it appears that the classification of subsamples with an log (LR) ≥ 1 by PG into separate putative contributor clusters that are then suitable for PG replicate analysis is a valid approach at this time. Other subsample classification systems based upon genetic distance considerations are currently under investigation.

In a single mixture, the DSCS procedure produces up to six different subsample types with associated LRs from separate putative contributors. These subsample types include 1- and 2-cell subsamples, 2-cell mini-mixture subsamples, 1- and 2-cell replicates, and mini-mixture replicates. After replicate analysis, the maximum replicate likelihood ratio obtained is reported.

While the results are encouraging, more work is required before the DSCS method would be suitable for casework applications. The data set is limited, with 5 mixtures (2-6 persons) being analyzed in detail by DSCS, although approximately 300 separate STR genotyping reactions were performed on the mixture subsamples (40-80 per complex mixture) and several hundred more for the PG validation studies. Thus, a wider range of complex mixtures with differing mixture contributor ratios as well as environmentally compromised cells and different cell types (including vaginal epithelia, white blood cells and cells from touch DNA deposits) should be investigated to determine whether a similar improvement in contributor genotype information recovery compared to standard bulk mixture analysis is achieved.

It should be emphasized that DSCS, even if subsequent studies confirm its accuracy and value with *bona fide* forensic samples, is not intended to replace current standard methods of mixture analysis. Instead, it is intended to be used for complex mixtures only <u>after</u> a standard (‘bulk’) mixture analysis is performed, which results in a determination of the minimum number of contributors and the computed LRs for the POIs. Only then, a determination would be made as to whether DSCS would be worthwhile pursuing in an attempt to retrieve more probative information from the sample. Of course, an appropriate sampling strategy would have to be developed, validated and employed to permit subsequent DSCS analysis after standard analysis (without compromising the latter). For example, loosely adhering deposited cells on the underlying substrate could be transferred directly [17] or indirectly (via a cotton swab) onto the inert silicone Gel-film™ before or after standard bulk mixture analysis. DSCS’s main use would be retrospective for those more complex mixture cases in which one (or more) POI contributor(s) has returned an LR that did not provide “extremely strong support” for the inclusionary hypothesis (e.g., did not reach the internationally recognized threshold of a log (LR) > 6, beyond which the verbal conclusion does not change [29,30]) or in cases where at least one POI was not present in a high enough relative proportion for standard DNA mixture analysis methods to retrieve an STR profile. Also, the method could prove to be particularly useful for those complex mixture cases in which multiple and/or alternative first-degree relatives are the POI(s).

Since the DSCS method uses morphologically distinguishable features to recognize cells for recovery, a potential concern would be whether using the method in real forensic physiological dried stains might therefore fail to pick up all DNA contributors to the mixture due to the presence of cell free DNA (cfDNA) [36] which would be missed if only cells were collected from the sample. The authors do not believe that the presence of cfDNA *per se* invalidates the proposed DSCS approach. Even if there were cfDNA contributors to a mixture in the absence of concomitant deposited cells, their presence would be detected by standard bulk analysis (assuming the cfDNA was of sufficient quality and quantity for STR analysis, otherwise its presence would be moot). So, in these type of mixtures DSCS would fail to detect the cfDNA contributor(s) and improved genotype information recovery from that individual (or individuals) by DSCS would not be possible. However, the ability to recover more genotype information would remain the same for any other contributors to the same mixture that did have cells present, including from those that may not have been detectable by standard methods due to a very low mixture ratio. Indeed, arguably there would now be a higher probability of recovering genotype information by DSCS from a minor sub-analytical contributor since the mixture ratio of this minor donor in the cellular fraction would now be increased compared to that in the bulk mixture (a proportion of which would comprise cfDNA from a non-cellular donor).

The question arises as to whether the results obtained from DSCS would have limited utility since they cannot be entered into searchable crime scene and criminal databases such as CODIS. Currently, results of standard complex mixture analysis cannot be input into CODIS either. This is because deconvolution into the constituent genotypes with these mixtures using PG is imprecise in that for every locus there are usually several genotype combinations that could feasibly (albeit with different probabilities) explain the results. An exception to this would be where there is a clear and easily resolvable major donor, but unfortunately, even in these cases, the POI may not be the major donor. Very often the most probable multi-locus genotype inferred by the PG system is not exactly that of the POI. This uncertainty of the genotype obtained by PG currently precludes the ability to input a particular source genotype into CODIS. Notwithstanding the above, there are ongoing efforts to produce software to enable standard PG mixture analysis results to search CODIS type databases in the future (e.g. STRmix’s DBLR^TM^ investigative tool). The latter envisions the possibility of a local search of the crime scene mixture by interrogating individuals from the criminal database via a rapid calculation of LRs from each of the individuals (or subsets thereof). However, with the DSCS method, single source genotypes can be obtained and, arguably, there would actually be a better chance of being able to interrogate donors of complex mixtures against CODIS than the current method. This scenario however would have to be accompanied by extensive community validation of the low template methods employed by DSCS which admittedly would require extra work on the part of the broader community. Many labs are, in reality, using low template methods currently due to the use of high sensitivity kits and the fact that PG deals routinely with the stochastic effects common to low template samples [37]. The path to CODIS or other regulated database acceptability may be long and narrow for DSCS approaches employing high sensitivity methods, but it is not impossible, since there are published guidelines on how to validate them [38].

If a community developmental validation of DSCS with more casework relevant samples proved successful, DSCS using simplified micromanipulation as used here could readily be implemented in a casework laboratory due to it only requiring basic equipment and materials and its ease of use. Alternatively, the use of a semi-automated digital cell sorting method [18] could be employed for those labs that prefer an instrument-versus microscope-based approach. The PG software used in this study is perhaps already being used or validated in the laboratory.

In summary, the study establishes a method that could potentially overcome limitations to the current DNA mixture analysis paradigm. The physical sampling with PG analysis of captured cells permits the recovery of most (or all) of the contributor STR genotype information contained within a complex mixture. Indeed, low level contributors that may not even be detectable by standard methods could benefit from a subsampling method such as proposed here to retrieve the otherwise totally lost information. Eventual implementation of such a DSCS approach would be expected to increase the number of complex DNA mixture cases in which a person of interest is definitively implicated or excluded as being a contributor to the biological trace.

## Declaration of interest

No competing interests to disclose.

## Funding Source

The authors would like to thank the State of Florida for initial seed funding for this project. The funders had no role in study design; in the collection, analysis, and interpretation of data; in the writing of the report; and in the decision to submit the article for publication.

## Abbreviations

DSCS: (direct single cell subsampling);
POI: (person-of-interest);
PG: (probabilistic genotyping);
NOC: (number of contributors)

## Acknowledgments

The authors would like to thank the anonymous donors who provided samples for this study and STRmix™ Technical and Scientific Support for assistance answering technical questions related to the PG system as well as Catherine McGovern (Environmental Science and Research) for assistance in running the 5-person bulk mixture and the 6-person bulk mixture with STRmix™ v2.9.

## Notes

### Competing Interest Statement

The authors have declared no competing interest.

