## Supplementary Figures and Tables for "Probabilistic Genotyping of Single Cell Replicates from Complex DNA Mixtures Recovers Higher Contributor LRs than Standard Analysis"

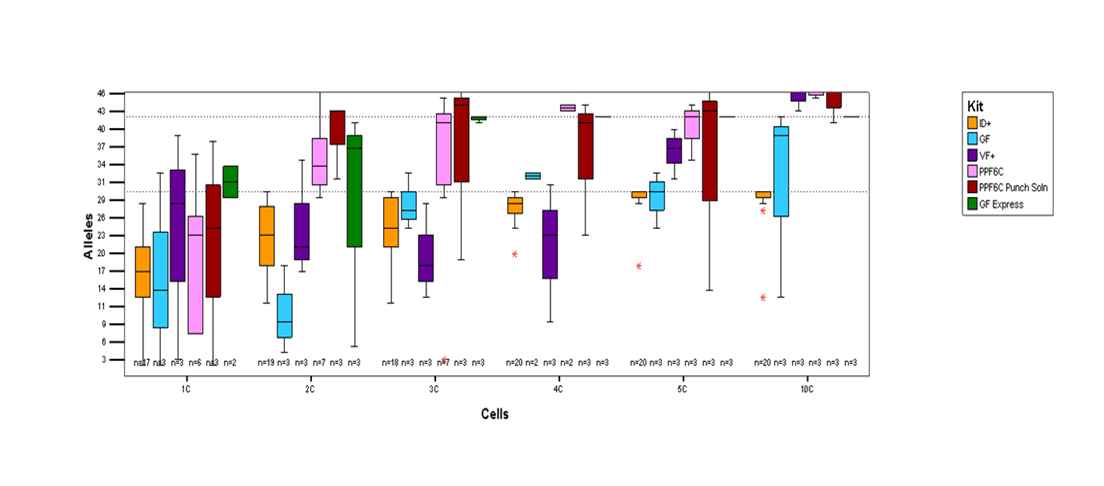

**Supp Figure 1.** **STR kits/lysis conditions.** 1-10 cell subsamples were collected from single source buccal slides using various STR kits and lysis conditions. Identifiler^®^ Plus (orange), GlobalFiler^®^ (light blue), VeriFiler^®^ Plus (purple), PowerPlex^®^ Fusion 6C (pink), PowerPlex^®^ Fusion 6C + amp solution and lysed with punch solution (burgundy), and GlobalFiler^®^ Express (green). The lower dotted line indicates a full profile for Identifiler^®^ Plus while the higher dotted line indicates a full profile for GlobalFiler^®^ systems.

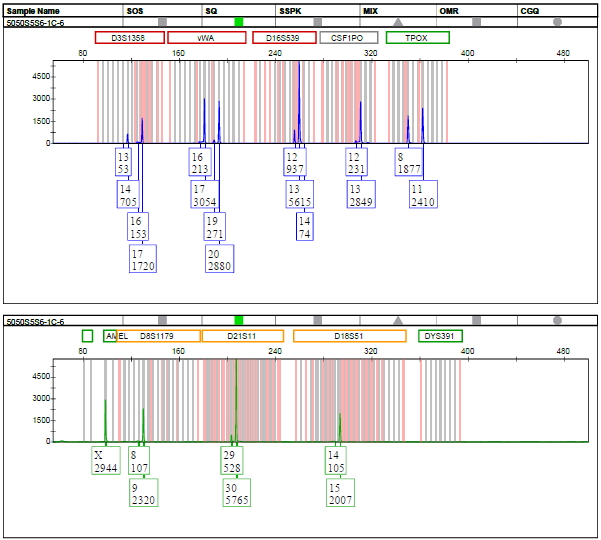

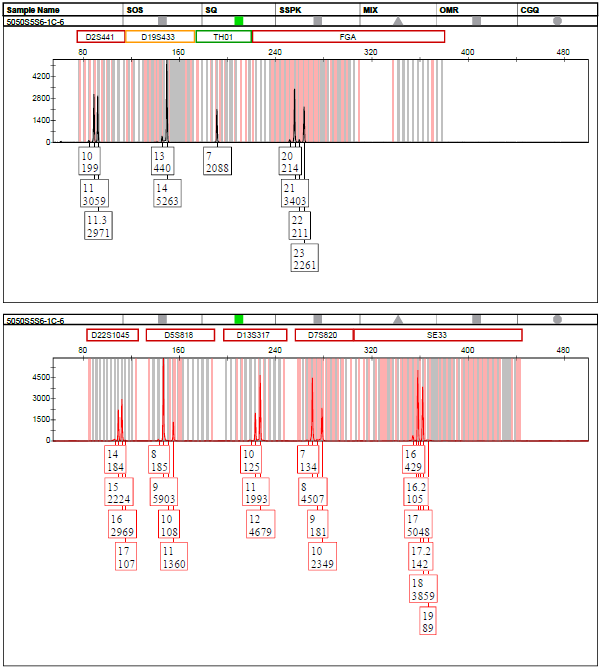

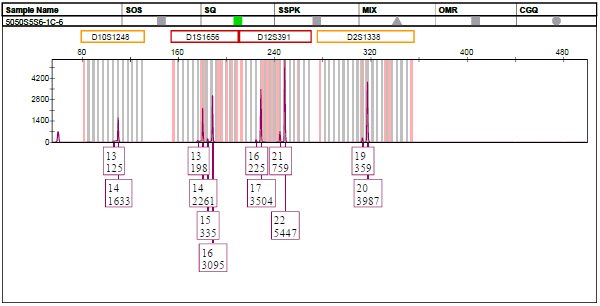

**Supp. Figure 2. Partial single source profile (1-cell)**

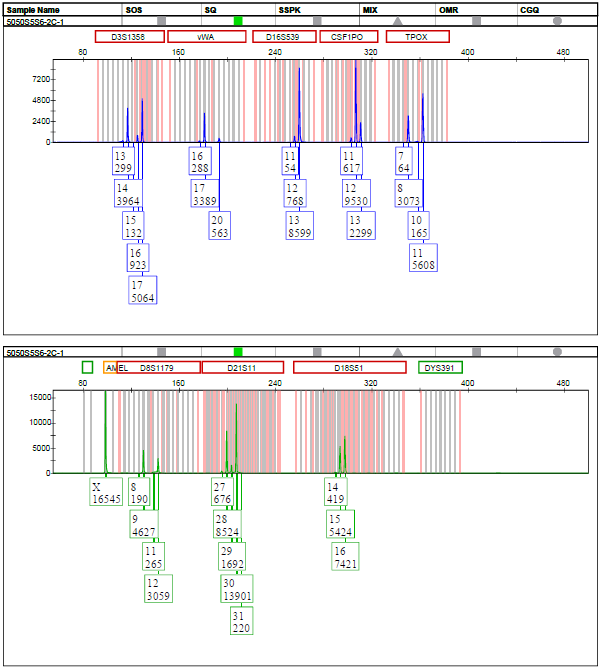

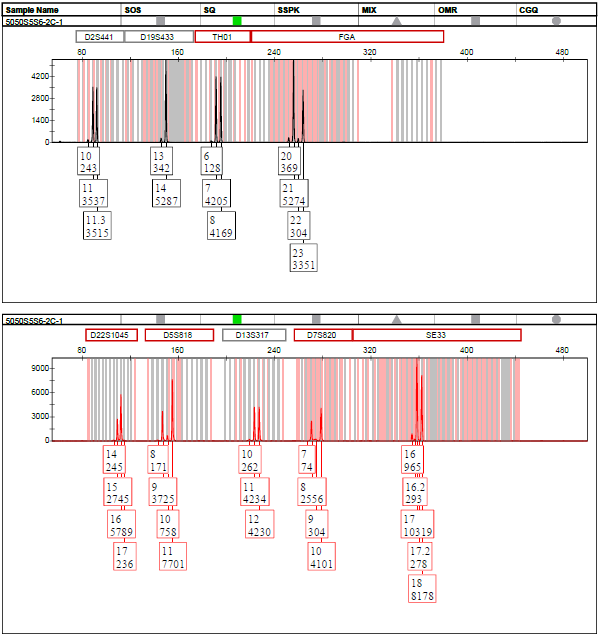

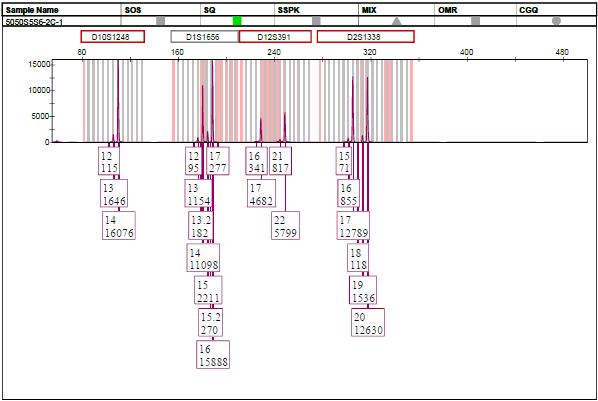

**Supp. Figure 3. Full single source profile (2-cells)**

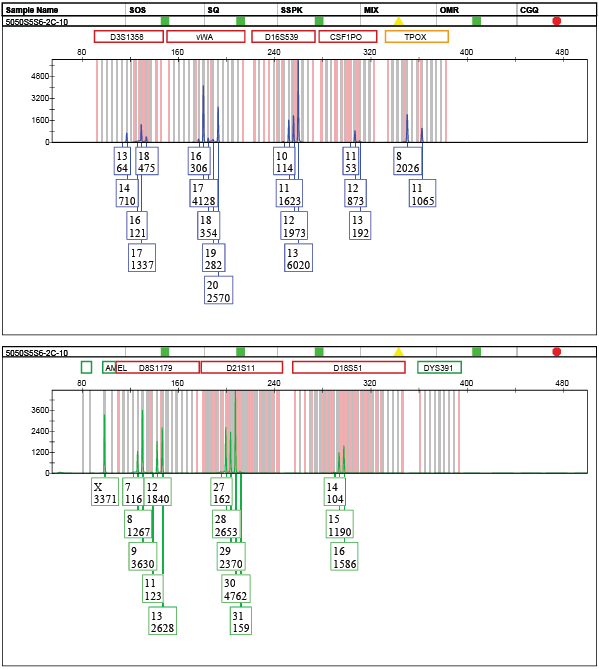

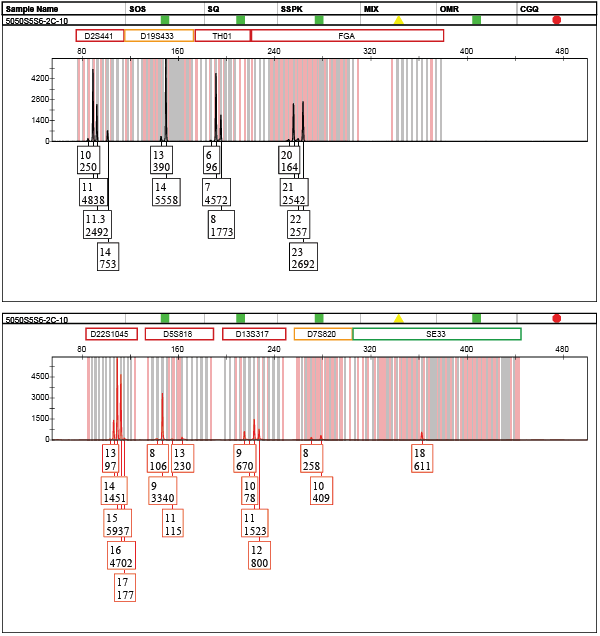

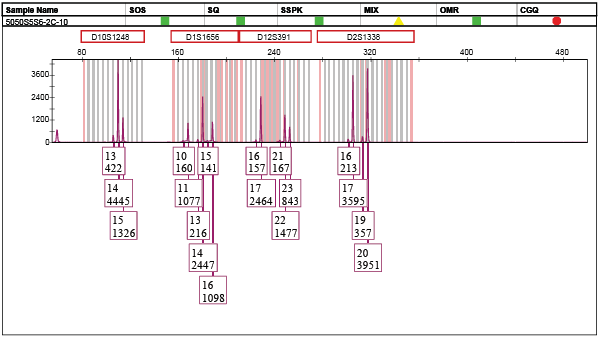

**Supp. Figure 4. Mixed profile (2-cells)**

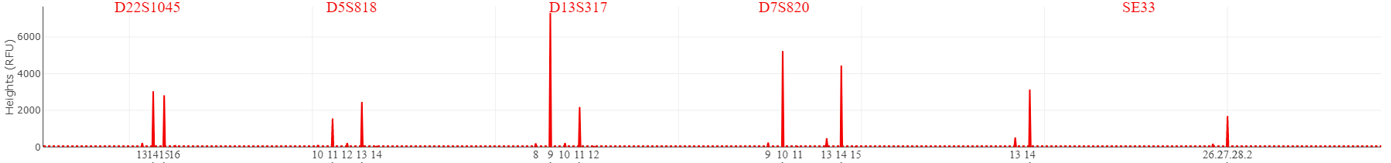

B

A

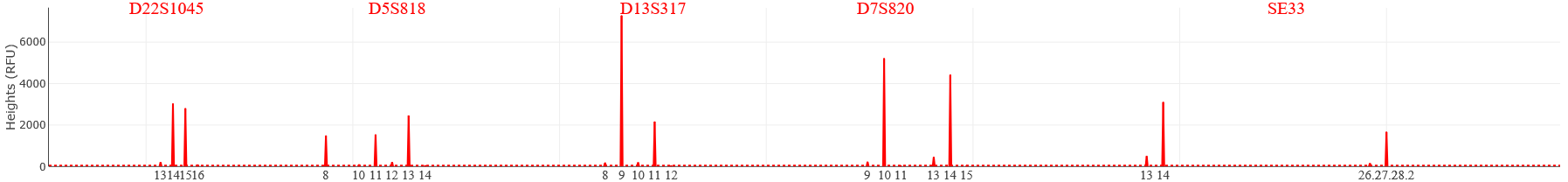

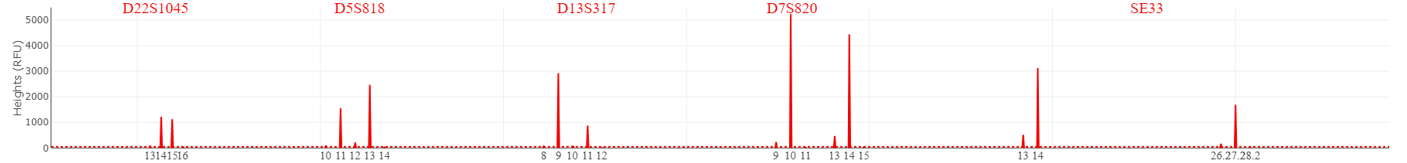

C

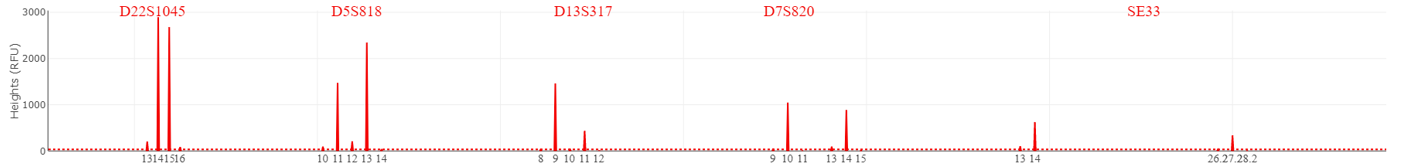

D

**Supp Figure 5. PG DSCS Parameters Test.** Regular sample (A), Sample with drop-in (B), Inhibited sample (C), Degraded sample (D)

| Non-shared Alleles | Known Contributor |
| --- | --- |
| 17 | 14 |
| 3 | -3 |
| 23 | 17 |
| 12 | -7 |
| 15 | -2 |
| 6 | 7 |
| 20 | 8 |
| 17 | 6 |
| 10 | 5 |
| 4 | 0 |
| 14 | -1 |
| 10 | -2 |
| 16 | -1 |
| 6 | -1 |
| 16 | 19 |
| 6 | -8 |
| 10 | 3 |
| 4 | -1 |
| 16 | 15 |
| 9 | -6 |
| 16 | 17 |
| 8 | -5 |
| 9 | 8 |
| 2 | -2 |
| 4 | 1 |
| 11 | 4 |
| 14 | 12 |
| 13 | -20 |
| 3 | 4 |
| 4 | -3 |
| 7 | -1 |
| 13 | 4 |
| 8 | 11 |
| 4 | 8 |
| 1 | -6 |
| 1 | 0 |
| 8 | 4 |
| 6 | -1 |
| 1 | -2 |
| 6 | 2 |
| 6 | 10 |
| 6 | 11 |
| 3 | -3 |
| 2 | 0 |
| 1 | -2 |
| 3 | -1 |
| 1 | 1 |
| 5 | -4 |
| 2 | -1 |
| 2 | 2 |
| 1 | -1 |
| 4 | 14 |
| 3 | -7 |
| 7 | 14 |
| 1 | -2 |
| 4 | 5 |
| 6 | 7 |
| 3 | 2 |
| 3 | 0 |
| 4 | 4 |
| 2 | -1 |
| 2 | 2 |
| 1 | -3 |
| 3 | 4 |
| 1 | -2 |
| 1 | 0 |
| 0 | -1 |
| 6 | 12 |
| 7 | 0 |
| 3 | 4 |
| 1 | -1 |
| 3 | 0 |
| 1 | -1 |
| 3 | 1 |
| 4 | 0 |
| 7 | 10 |
| 7 | 17 |
| 4 | 1 |
| 6 | 16 |
| 20 | 9 |
| 1 | -1 |
| 1 | 0 |
| 2 | 5 |
| 5 | 9 |
| 2 | -3 |
| 4 | 0 |
| 1 | 0 |
| 2 | 6 |
| 1 | -1 |
| 1 | 3 |
| 7 | 6 |
| 6 | 15 |
| 0 | -1 |
| 2 | 1 |
| 0 | 3 |
| 1 | -2 |
| 1 | 3 |
| 6 | 10 |
| 6 | 11 |
| 4 | 0 |
| 0 | -1 |

| Non-shared Alleles | Non-contributor |
| --- | --- |
| 3 | 1 |
| 3 | 0 |
| 3 | -1 |
| 3 | -2 |
| 3 | -3 |
| 3 | -4 |
| 3 | -5 |
| 12 | -8 |
| 12 | -9 |
| 12 | -11 |
| 12 | -12 |
| 12 | -13 |
| 12 | -14 |
| 12 | -15 |
| 12 | -16 |
| 12 | -17 |
| 12 | -18 |
| 12 | -19 |
| 12 | -20 |
| 12 | -21 |
| 12 | -22 |
| 12 | -23 |
| 12 | -24 |
| 12 | -25 |
| 12 | -26 |
| 12 | -27 |
| 12 | -28 |
| 12 | -29 |
| 12 | -30 |
| 12 | -31 |
| 12 | -32 |
| 12 | -33 |
| 12 | -34 |
| 12 | -35 |
| 12 | -36 |
| 12 | -37 |
| 12 | -38 |
| 12 | -39 |
| 12 | -40 |
| 12 | -41 |
| 12 | -44 |
| 12 | -50 |
| 6 | 2 |
| 6 | 1 |
| 6 | 0 |
| 6 | -1 |
| 6 | -2 |
| 6 | -3 |
| 6 | -4 |
| 6 | -5 |
| 6 | -6 |
| 6 | -7 |
| 17 | -5 |
| 17 | -7 |
| 17 | -8 |
| 17 | -9 |
| 17 | -10 |
| 17 | -11 |
| 17 | -12 |
| 17 | -13 |
| 17 | -14 |
| 17 | -15 |
| 17 | -16 |
| 17 | -17 |
| 17 | -18 |
| 17 | -19 |
| 17 | -20 |
| 17 | -21 |
| 17 | -22 |
| 17 | -23 |
| 17 | -24 |
| 17 | -25 |
| 17 | -26 |
| 17 | -27 |
| 17 | -28 |
| 17 | -29 |
| 17 | -31 |
| 17 | -35 |
| 17 | -36 |
| 17 | -50 |
| 4 | 2 |
| 4 | 1 |
| 4 | 0 |
| 4 | -1 |
| 4 | -2 |
| 4 | -3 |
| 4 | -4 |
| 10 | -50 |
| 10 | -1 |
| 10 | -2 |
| 10 | -3 |
| 10 | -4 |
| 10 | -5 |
| 10 | -6 |
| 10 | -7 |
| 10 | -8 |
| 10 | -9 |
| 10 | -10 |
| 10 | -11 |
| 10 | -12 |
| 10 | -13 |
| 10 | -14 |
| 10 | -15 |
| 10 | -16 |
| 10 | -17 |
| 10 | -18 |
| 10 | -19 |
| 10 | -20 |
| 6 | 1 |
| 6 | 0 |
| 6 | -1 |
| 6 | -2 |
| 6 | -3 |
| 6 | -4 |
| 6 | -5 |
| 6 | -50 |
| 6 | 0 |
| 6 | -1 |
| 6 | -2 |
| 6 | -3 |
| 6 | -4 |
| 6 | -5 |
| 6 | -6 |
| 6 | -7 |
| 6 | -8 |
| 6 | -9 |
| 6 | -10 |
| 6 | -11 |
| 6 | -12 |
| 4 | 1 |
| 4 | 0 |
| 4 | -1 |
| 4 | -2 |
| 4 | -3 |
| 4 | -4 |
| 9 | 0 |
| 9 | -2 |
| 9 | -3 |
| 9 | -4 |
| 9 | -5 |
| 9 | -6 |
| 9 | -7 |
| 9 | -8 |
| 9 | -9 |
| 9 | -10 |
| 9 | -11 |
| 9 | -12 |
| 9 | -13 |
| 9 | -14 |
| 9 | -15 |
| 9 | -16 |
| 9 | -17 |
| 8 | -50 |
| 8 | -1 |
| 8 | -2 |
| 8 | -3 |
| 8 | -4 |
| 8 | -5 |
| 8 | -6 |
| 8 | -7 |
| 8 | -8 |
| 8 | -9 |
| 8 | -10 |
| 8 | -11 |
| 8 | -12 |
| 8 | -13 |
| 8 | -14 |
| 8 | -15 |
| 8 | -16 |
| 8 | -17 |
| 8 | -19 |
| 8 | -20 |
| 8 | -21 |
| 8 | -22 |
| 8 | -23 |
| 8 | -24 |
| 8 | -25 |
| 8 | -26 |
| 8 | -27 |
| 8 | -28 |
| 9 | 2 |
| 2 | 1 |
| 2 | 0 |
| 2 | -1 |
| 2 | -2 |
| 2 | -3 |
| 2 | -4 |
| 2 | -5 |
| 2 | -6 |

| Non-shared Alleles | log(LR) |
| --- | --- |
| -3 | 6 |
| 45 | 6 |

**Supp Figure 6.** **Specificity of DSCS (STRmix^TM^) mini-mixture analysis.** Single source subsamples. LR = 0 plotted as -50. Blue circles (known contributors), orange circles (non-contributor)

| log(LR) | Known Contributors |
| --- | --- |
| 7.28604 | 17.9035 |

| log(LR) | Non-Contributors |
| --- | --- |
| -2.04359 | -1.56827 |
| -4.31999 | -2.3451 |
| -1.44605 | -0.759157 |
| -0.642951 | -1.44254 |
| -2.3717 | -1.1683 |
| -3.15456 | -1.30609 |
| -0.786762 | -5.71785 |
| -2.93104 | -2.86156 |
| -0.762728 | -2.21579 |
| -0.00226462 | -0.192603 |
| -1.60281 | -1.8408 |
| -3.01384 | -1.33015 |
| -0.736888 | -0.712765 |
| -4.35148 | -7.09012 |
| -0.481068 | 0.423806 |
| -1.43934 | -2.59969 |
| 2.43612 | 0.769612 |
| 0.0937168 | -0.893372 |
| -1.89762 | -3.02145 |
| -1.14827 | -2.51117 |
| 0.775312 | -0.739955 |
| -0.862166 | -1.24998 |
| -0.117732 | 0.207185 |
| -1.36698 | -1.99864 |
| -1.30885 | -3.05045 |
| -1.68897 | -6.30975 |
| -2.21627 | -4.26367 |
| -0.687438 | -1.28234 |
| -1.49271 | -7.99535 |
| -2.53704 | -3.10124 |
| -1.6554 | -3.37451 |
| -0.543449 | -1.75469 |
| -0.615334 | -1.19005 |
| -0.650609 | -3.30298 |
| -0.107846 | -2.60257 |
| -0.40128 | -7.81552 |
| -0.0985752 | -1.53934 |
| -1.29623 | -1.91378 |
| -0.808263 | -2.41371 |
| -0.994341 | -0.855897 |
| -4.46884 | -2.4386 |
| -1.84044 | -0.504714 |
| -1.36951 | -2.47881 |
| -0.358828 | -0.548771 |
| -4.66219 | -1.74312 |
| -2.22295 | -2.83494 |
| -4.11428 | -1.89819 |
| 0.0358225 | -1.32649 |
| -1.5904 | -3.00564 |
| -1.90212 | -1.79905 |
| 0.460583 | -1.48882 |
| -3.36631 | -1.67723 |
| -1.01164 | -1.73836 |
| -2.41496 | -4.6062 |
| -0.662343 | -0.878167 |
| -1.71378 | -3.23297 |
| -1.38885 | -3.6149 |
| -2.70347 | -2.12941 |
| -1.20383 | -6.52445 |
| -3.63533 | -3.05778 |
| -2.70094 | -0.877501 |
| -0.834455 | -0.466054 |
| -1.15652 | -2.62483 |
| -0.623283 | -0.197851 |
| -4.21916 | -2.29491 |
| -2.70222 | -2.41966 |
| -2.97342 | -0.363252 |
| -2.28244 | -1.4616 |
| -0.923149 | -2.95871 |
| -1.9786 | -2.02323 |
| -4.51746 | -1.94416 |
| -2.32162 | -0.473404 |
| 0.0390466 | -2.82402 |
| -2.01989 | -2.09358 |
| -1.50472 | -1.93982 |
| -2.54392 | -2.89475 |
| -0.8536 | -0.0333365 |
| 0.256476 | -0.607985 |
| -2.53691 | -0.891345 |
| -1.82537 | -1.06786 |
| -2.12308 | -1.42552 |
| -2.17141 | -1.11616 |
| -4.52697 | -3.08716 |
| -1.2691 | -3.10875 |
| -2.59414 | -1.18756 |
| -3.10661 | -2.57626 |
| -3.24616 | -0.454788 |
| -1.00772 | -2.11837 |
| -2.89023 | -3.58124 |
| -0.992242 | -1.66975 |
| 0.199567 | -6.83207 |
| -1.44573 | -3.07195 |
| -2.14841 | -6.86373 |
| 0.0862861 | -0.978986 |
| -0.013279 | -0.742231 |
| -2.15605 | -5.97262 |
| -1.1877 | -0.785516 |
| -1.71421 | -1.93096 |
| -4.43661 | -3.45554 |
| -3.04303 | -5.17703 |
| -1.37049 | -2.73016 |
| -3.17157 | -0.530185 |
| -2.10805 | -6.19185 |
| -2.22771 | -1.00706 |
| -3.10017 | -2.54568 |
| 0.115088 | -1.74078 |
| -2.63878 | -2.75564 |
| -4.48063 | -3.29919 |
| -2.91108 | -2.33516 |
| -2.82668 | -2.34201 |
| -0.75549 | -2.57805 |
| -0.918364 | -4.71132 |
| -4.54622 | -1.6273 |
| -4.15243 | -2.16985 |
| -0.417182 | -0.316378 |
| -3.65229 | -2.26414 |
| 1.04512 | 0.809076 |
| -4.79877 | -2.8689 |
| -0.48564 | -1.28678 |
| -0.451271 | -2.68679 |
| -4.01547 | -4.28375 |
| -1.24763 | -6.34586 |
| -2.61852 | -0.405645 |
| -3.01975 | -7.51218 |
| -1.35443 | -2.18027 |
| -1.79374 | -1.8252 |
| -4.37357 | -2.8704 |
| -2.29121 | -8.45414 |
| -2.38018 | -2.62834 |
| -2.57979 | -2.66379 |
| -3.47969 | -1.46687 |
| -1.48723 | -2.41609 |
| -1.89251 | -2.7466 |
| -0.450743 | -6.16301 |
| -0.757263 | -0.828034 |
| -1.791 | -2.88079 |
| -1.00903 | 0.256673 |
| -3.60478 | -1.52856 |
| -1.08834 | -0.954882 |
| -2.65603 | -1.07009 |
| -1.63218 | -0.744032 |
| -2.82501 | -3.35151 |
| -0.986254 | -1.61887 |
| -2.63258 | -2.99274 |
| -1.12678 | -1.82054 |
| -1.30441 | -0.439097 |
| -0.138778 | -2.84202 |
| 0.059044 | 0.445901 |
| -1.19168 | -5.46995 |
| -2.54246 | -1.75973 |
| -2.44707 | 1.03731 |
| -0.585127 | -1.14744 |
| -3.26159 | -6.35532 |
| -4.72882 | -3.29672 |
| -3.17345 | -1.4605 |
| -3.67165 | -3.67995 |
| -2.29851 | -2.12515 |
| -2.14191 | -2.29138 |
| -2.11943 | -2.99808 |
| -1.06905 | -1.9026 |
| -1.59445 | -1.86382 |
| -3.86456 | -2.69084 |
| -2.79806 | -0.737426 |
| -3.16571 | -1.96958 |
| -1.75062 | -2.85316 |
| 1.23456 | -2.2377 |
| -0.670397 | -3.25243 |
| -2.98494 | -0.359069 |
| -3.77581 | -7.12207 |
| -2.7983 | -2.67832 |
| -1.18127 | -1.34834 |
| -1.75366 | -0.725234 |
| -4.13716 | -1.59156 |
| -0.158214 | -0.660582 |
| -0.872053 | -1.08659 |
| -2.69691 | -3.0209 |
| -1.46783 | -3.49758 |
| -3.69196 | -8.59531 |
| -1.90971 | -5.7538 |
| -3.05385 | -1.78607 |
| -0.036768 | 0.830148 |
| -5.66324 | -2.79239 |
| -2.47559 | -0.663515 |
| 0.0985421 | 0.465945 |
| -2.02369 | -0.833683 |
| -0.774914 | -0.0825005 |
| -1.30648 | -3.39599 |
| -1.14404 | -0.904019 |
| -2.8209 | -2.98377 |
| -0.0893965 | -2.78254 |
| -1.31086 | -1.78801 |
| -1.44768 | -2.60437 |
| -0.969359 | -0.252095 |
| -0.273634 | -0.357471 |
| -2.04139 | -1.07258 |
| -2.06666 | -0.193342 |
| -3.44746 | -2.21934 |
| -1.89634 | -7.07427 |
| -1.02224 | -1.08583 |
| -2.65465 | -8.33695 |

| log(LR) | log(LR)replicates |
| --- | --- |
| -60 | -60 |
| 60 | 60 |

**Supp Figure 7.** **Specificity improvement of DSCS (STRmix^TM^) with mini-mixture replicate analysis*.*** Known contributors above the diagonal (i.e. y=x) represent improved LR recovery from replicate analysis. Single Source Cell Replicates. LR = 0 plotted as -50. Blue circles (known contributors); orange circles (non-contributors)

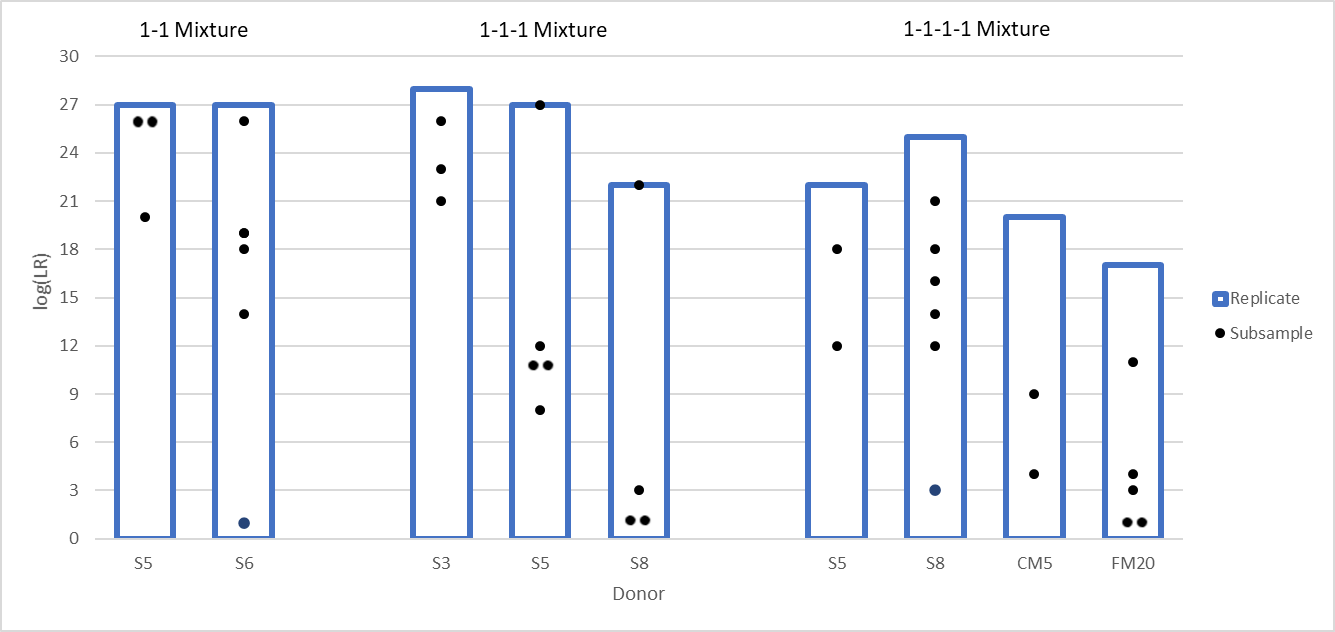

**Supp Figure 8. STRmix 2, 3, and 4-person mixture replicate log(LR) values with the respective subsamples used for replicate analysis.**

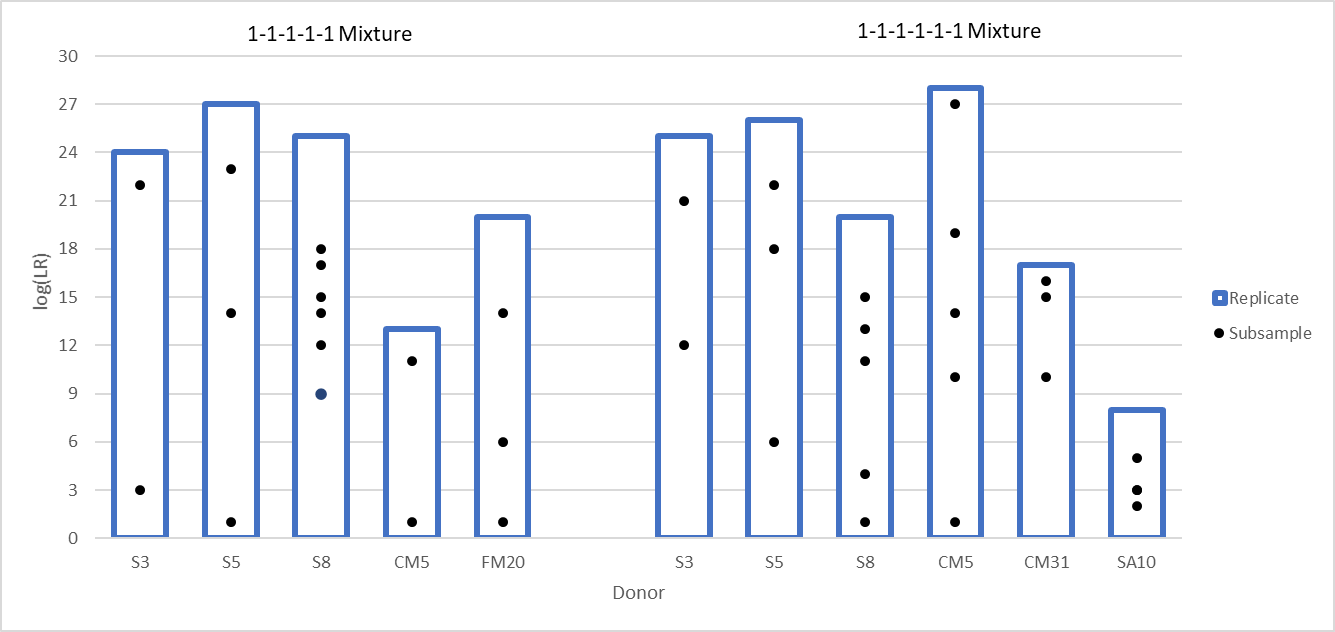
**Supp Figure 9. STRmix 5 and 6-person replicate log(LR) values with the respective subsamples used for replicate analysis.**

log(LR)

| Donor | Reference | DSCS Replicate |
| --- | --- | --- |
| S3 | 28 | 19 |
| S6.2 | 27 | 14 |
| SA70 | 32 | 19 |
| SA79 | 25 | 24 |
| SA80 | 28 | 20 |
| SA10 | 25 | 13 |

**Supp Figure 10. Contributor log (LR) recovery by DSCS (STRmix^TM^) in an additional 6-person mixture**

**Supplementary Tables**

**Supp. Table 1. STRmix^TM^ DSCS Stutter Parameters**

| Parameter | α | β | MODE | Max Stutter % |
| --- | --- | --- | --- | --- |
| Allele Variance c^2^ | 26.546 | 8.638 | 220.666 | NA |
| Back Stutter Variance k^2^ | 1.789 | 58.577 | 46.217 | 0.7 |
| Forward Stutter Variance k^2^ | 1.623 | 30.254 | 18.848 | 0.7 |
| Double Back Stutter Variance k^2^ | 1.501 | 16.203 | 8.118 | 0.3 |
| -2bp Stutter Variance k^2^ | 5.753 | 7.883 | 37.468 | 0.5 |
| +2bp Stutter Variance k^2^ | 11.679 | 15.925 | 170.063 | 0.15 |
| -6bp Stutter Variance k^2^ | 7.923 | 10.372 | 71.805 | 0.15 |
| LSAE Variance | 0.112 | | | |

**Supp. Table 2. STRmix^TM^ Bulk Mixture Stutter Parameters**

| Parameter | α | β | MODE | Max Stutter % |
| --- | --- | --- | --- | --- |
| Allele Variance c^2^ | 8.902 | 5.808 | 45.895 | NA |
| Back Stutter Variance k^2^ | 1.5 | 25.849 | 12.925 | 0.7 |
| Forward Stutter Variance k^2^ | 1.655 | 9.338 | 6.116 | 0.15 |
| Double Back Stutter Variance k^2^ | 6.756 | 2.081 | 11.978 | 0.05 |
| -2bp Stutter Variance k^2^ | 1.988 | 10.832 | 10.702 | 0.1 |
| +2bp Stutter Variance k^2^ | NA | NA | NA | NA |
| -6bp Stutter Variance k^2^ | NA | NA | NA | NA |
| LSAE Variance | 0.032 | | | |

**Supp. Table 3. Additional STRmix^TM^ Settings**

| **Method** | **Burn-in Accepts** | **Post Burn-in Accepts** | **Degradation Max** | **Saturation Threshold (RFUs)** |
| --- | --- | --- | --- | --- |
| DSCS | 100,000 | 500,000 | 0.1 | 30,000 |
| Bulk Mixture | 10,000 | 50,000 | 0.01 | 30,000 |

**Supp. Table 4. Test DSCS Samples**

| Sample | STRmix^TM^ log(LR) |
| --- | --- |
| S5-3C-1 | 27 |
| S5-3C-1 drop-in | 27 |
| S5-3C-1 inhibited | 27 |
| S5-3C-1 degraded | 27 |

**Supp. Table 5. Percentage of subsamples and replicates analyzed with EuroForMix that reach the specified threshold**

| Sample Type | Subsample | Replicate | Subsample | Replicate |
| --- | --- | --- | --- | --- |
|  | log(LR)>0 | | log(LR)≥6 | |
| SS 1 cell (s=100) | 87% | 100% | 45% | 94% |
| SS 2 cell (s=81) | 89% | 100% | 64% | 93% |
| Mix 2 cell (s=110) | 60% | 100% | 30% | 93% |
